# Is brightfield all you need for mechanism of action prediction?

**DOI:** 10.1101/2022.10.12.511869

**Authors:** Ankit Gupta, Philip J Harrison, Håkan Wieslander, Jonne Rietdijk, Jordi Carreras Puigvert, Polina Georgiev, Carolina Wählby, Ola Spjuth, Ida-Maria Sintorn

**Affiliations:** Department of Information Technology and SciLifeLab, Uppsala University, Uppsala, Sweden; Department of Pharmaceutical Biosciences, Uppsala University, Uppsala, Sweden

**Author notes:** Equal contribution (Ankit Gupta, Philip J Harrison).

## Abstract

Fluorescence staining techniques, such as Cell Painting, together with fluorescence microscopy have proven invaluable for visualizing and quantifying the effects that drugs and other perturbations have on cultured cells. However, fluorescence microscopy is expensive, time-consuming, and labor-intensive, and the stains applied can be cytotoxic, interfering with the activity under study. The simplest form of microscopy, brightfield microscopy, lacks these downsides, but the images produced have low contrast and the cellular compartments are difficult to discern. Nevertheless, by harnessing deep learning, these brightfield images may still be sufficient for various predictive purposes. In this study, we compared the predictive performance of models trained on fluorescence images to those trained on brightfield images for predicting the mechanism of action (MoA) of different drugs. We also extracted CellProfiler features from the fluorescence images and used them to benchmark the performance. Overall, we found comparable and correlated predictive performance for the two imaging modalities. This is promising for future studies of MoAs in time-lapse experiments.

## 1 Introduction

Mechanism of action (MoA) describes the biological process by which a compound exhibits a pharmacological effect, such as the proteins targeted or the pathways modulated. Establishing a compound’s MoA provides particularly useful information for lead compounds prior to clinical trials and for identifying potentially adverse or toxic effects [1]. Various assays can be used to provide information on a compound’s MoA, including transcriptomics, proteomics and metabolomics assays [1]. Recently, morphology-based high-content imaging assays have proven beneficial for this task [2] and are also significantly easier and less expensive to scale to high-throughput than other assay types [3].

The simplest, cheapest, and least invasive form of light microscopy is brightfield (BF) microscopy. Due to the thinness and transparency of most cells, BF images typically have low contrast, making it difficult to detect internal cell structures. To overcome this limitation fluorescent dyes can be used. Fluorescence (FL) microscopy uses fluorescent dyes to stain specific targets (e.g. cell compartments) within the sample [4]. A noteworthy FL-based protocol is the Cell Painting assay [5] which combines six different stains to highlight eight different sub-cellular compartments. However, FL imaging is much more labor-intensive and expensive than BF imaging. The dyes required for imaging some cellular components can also be highly toxic to the cells. The problems become amplified for timelapse imaging experiments with many exposures [6]. Another more pervasive phototoxic effect in FL microscopy is photobleaching which not only decreases the fluorescent signal but also releases free-radicals [7].

Traditional analysis pipelines in image cytometry follow the path of identifying, segmenting, and extracting handcrafted quantitative features from the cells, often using the CellProfiler (CP) [8] software package. Common features include those related to size, shape, pixel intensity, and texture. For fluorescent images, these features can be extracted at the level of the various cellular compartments. These features can then be used as input to machine learning models. Alternatively, one can use deep learning methods [9], specifically convolutional neural networks (CNNs), to perform the predictive task directly from the raw pixel intensity data in an end-to-end data-driven fashion, circumventing the need for cell segmentation and determining which features to extract [10].

In this study, we compared the performance of CNNs trained on Cell Painting fluorescence images (five channels) to those trained on brightfield images (six z-planes) for predicting ten MoA classes for U2OS cells treated with various compounds. As a reference/benchmark, we also trained neural networks on CellProfiler-derived features from the fluorescence images. We explored the effect of including the DMSO solvent control data (wells with no compound treatment) as a predictive class in the models, as well as the influence of different normalization strategies. Example Cell Painting images and their BF counterparts for the ten MoA classes and the DMSO are shown in Figure 1.

**Figure 1:**
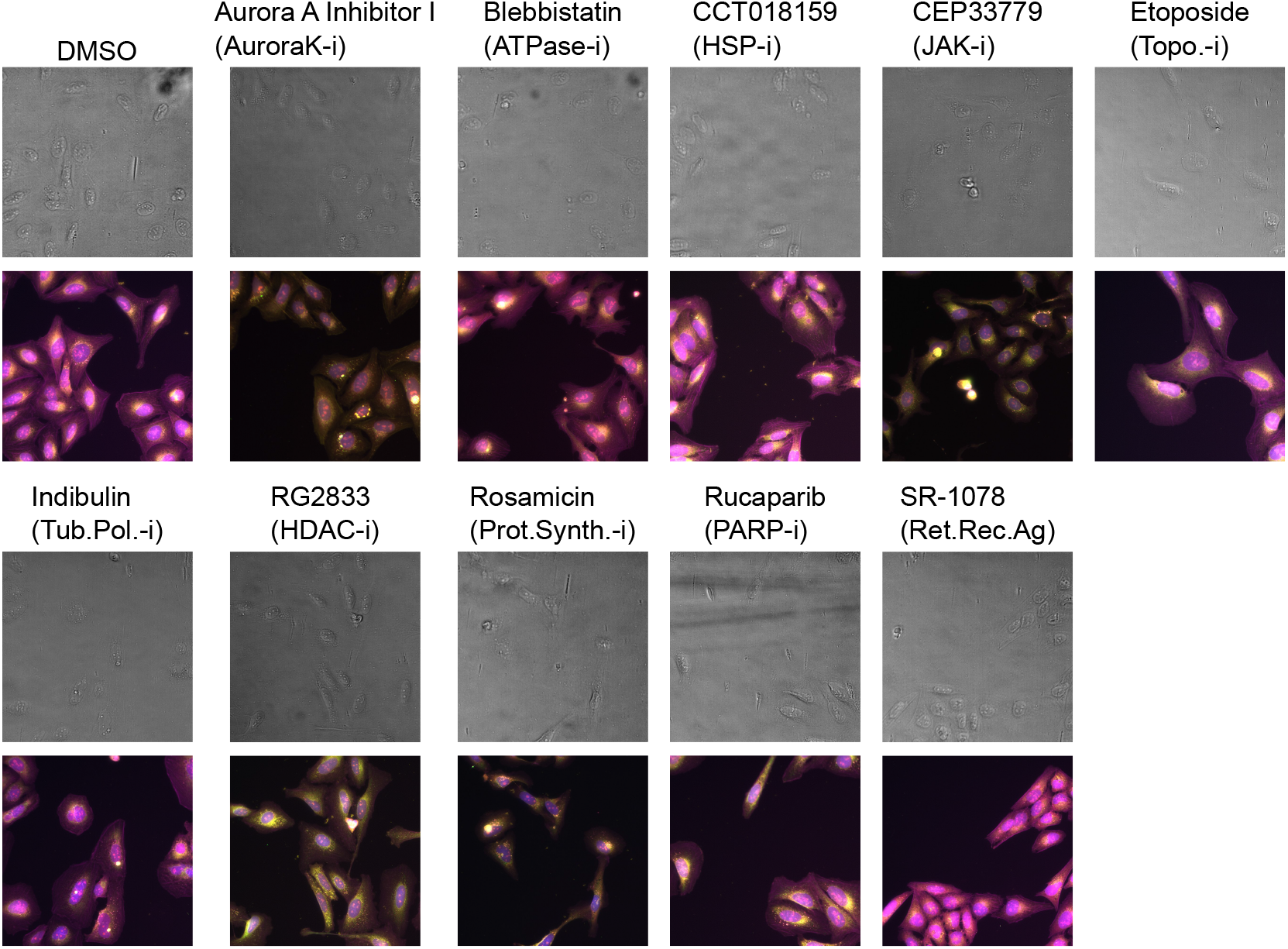
Example Cell Painting images for the ten MoA classes and the DMSO. The panel titles give the compound names for the selected images with the MoA abbreviation in parenthesis, where -i stands for inhibitor and Ag for agonist. The top images in each case show a maximum projection of the 6 BF z-planes, and those below show merged images for the FL channels with nuclei in blue, ER in green, RNA in magenta, Golgi/F-actin in red and mitochondria in yellow.

## 2 Methods

### Dataset

We used image data for compounds belonging to ten MoA classes (MoAs that we believed would be reasonably separable and that had a sufficient number of compounds (n) associated with them in our assay). The 10 MoAs were: ATPase inhibitors (ATPase-i, n = 18); Aurora kinase inhibitors (AuroraK-i, n = 20); HDAC inhibitors (HDAC-i, n = 33); HSP inhibitors (HSP-i, n = 24); JAK inhibitors (JAK-i, n = 21); PARP inhibitors (PARP-i, n = 21); protein synthesis inhibitors (Prot.Synth.-i, n = 23); retinoid receptor agonists (Ret.Rec.Ag, n = 19); topoisomerase inhibitors (Topo.-i, n = 32); and tubulin polymerization inhibitors (Tub.Pol.-i, n = 20). The compounds were administered at a dose of 10 *μ*M to U2OS cells, and exposed for 48 h, in 384 well plates. Each compound-level experiment was replicated six times. The compounds were distributed across eighteen microplates using PLAID (Plate Layouts using Artificial Intelligence Design, [11]), a constrained programming-based method that aims to limit unwanted bias and batch effects. Images (16-bit, 2160×2160 pixels) were captured with a 20X objective at five sites/fields-of-view in each well, with five fluorescence channels for the FL data and six evenly spaced z-planes for the BF data. See Supplementary Section 5.1 for more details.

### Data splitting

We performed five splits of the data, at the compound level, into training, validation, and test sets. The splitting was performed in a stratified manner based on the proportion of compounds for each MoA. Each of the five test sets contained approximately 20% of the data, with no overlap of compounds. For each split the remaining non-test data was shuffled, at the compound level, and assigned to training or validation in a stratified manner, with 80% to training and 20% to validation. DMSO data was added to the sets with five, one, and two wells per plate for training, validation, and testing, respectively.

### Model training

For all the comparisons made we used a standard ResNet-50 [12] model with consistent hyper-parameters and training strategies. See Supplementary Section 5.2 for the details.

### Normalization

The FL data was normalized using the mean and standard deviation of the pixel intensities of the control DMSO wells in each plate, to mitigate plate-level effects [13]. For BF, there is no well-established normalization protocol. In FL images, the information is present in both intensity and morphological variations. However, in BF images, only the morphological information is present, and thus, to establish a benchmark, different normalization strategies were explored (See Supplementary 5.3). The DMSO normalization turned out to be the best-performing normalization strategy also for BF.

### CellProfiler features

To benchmark the performance, we extracted CellProfiler (CP) features from the FL data on which we trained a fully-connected neural network with one hidden layer. We extracted 672 cell averaged features from the five channel FL image, including those related to (in CP parlance) ‘AreaShape’, ‘Correlation’, ‘Granularity’, ‘Intensity’, ‘Neighbors’, and ‘RadialDistribution’.

## 3 Result and Discussion

Table 1(a) shows comparative test set F1 scores for the five data splits (averaged across the MoA prediction classes) for the models trained on BF, FL, and CP data. Table 1(b) shows the combined F1 scores (across all five tests sets, i.e. for all the compounds in our dataset) with respect to the ten MoA classes and the DMSO. The BF models perform competitively with respect to the FL and CP models, the F1 scores are quite similar for all the MoAs, except for the retinoid receptor agonists (Ret.Rec.Ag), for which the BF results are slightly lagging. The most pronounced difference is for the DMSO class, for which the BF results are considerably lower than those for both FL and CP.

**Table 1:** Comparison of the results for the models trained on BF and FL images and CP features.

(a) Macro-F1 scores on the test sets for the five data splits for BF, FL and CP models.
| Modality | BF | FL | CP |
| --- | --- | --- | --- |
| Split 1 | 0.688 | **0.777** | 0.774 |
| Split 2 | 0.773 | 0.799 | **0.800** |
| Split 3 | 0.700 | **0.793** | 0.773 |
| Split 4 | 0.731 | **0.738** | 0.728 |
| Split 5 | 0.701 | 0.716 | **0.760** |

(b) F1 scores per MoA, across all five test sets, for BF, FL, and CP models.
| MoA | BF | FL | CP |
| --- | --- | --- | --- |
| ATPase-i | 0.674 | **0.701** | 0.699 |
| AuroraK-i | **0.742** | 0.675 | 0.677 |
| HDAC-i | **0.792** | 0.773 | 0.779 |
| HSP-i | 0.728 | **0.730** | 0.712 |
| JAK-i | 0.677 | 0.653 | **0.681** |
| PARP-i | 0.833 | 0.886 | **0.925** |
| Prot.Synth.-i | 0.768 | **0.793** | 0.770 |
| Ret.Rec.Ag | 0.684 | 0.769 | **0.795** |
| Topo.-i | 0.729 | 0.728 | **0.761** |
| Tub.Pol.-i | 0.881 | 0.854 | **0.887** |
| DMSO | 0.459 | **0.866** | 0.845 |
| Macro average | 0.724 | 0.766 | **0.776** |

### Feature Representation

We also visually explored the feature representation using UMAP projections for the BF and FL models compared to those based on CP features in Figure 2. As can be seen from Figure 2A., the DMSO samples (blue dots) are not clustered together but rather scattered among the MoAs. The same phenomenon can be seen in the FL models (Figure 2B.), although not as severely. This issue is also apparent in the reference CP features (Figure 2C.), where the DMSO samples are not well-separated from many of the MoA classes.

**Figure 2:**
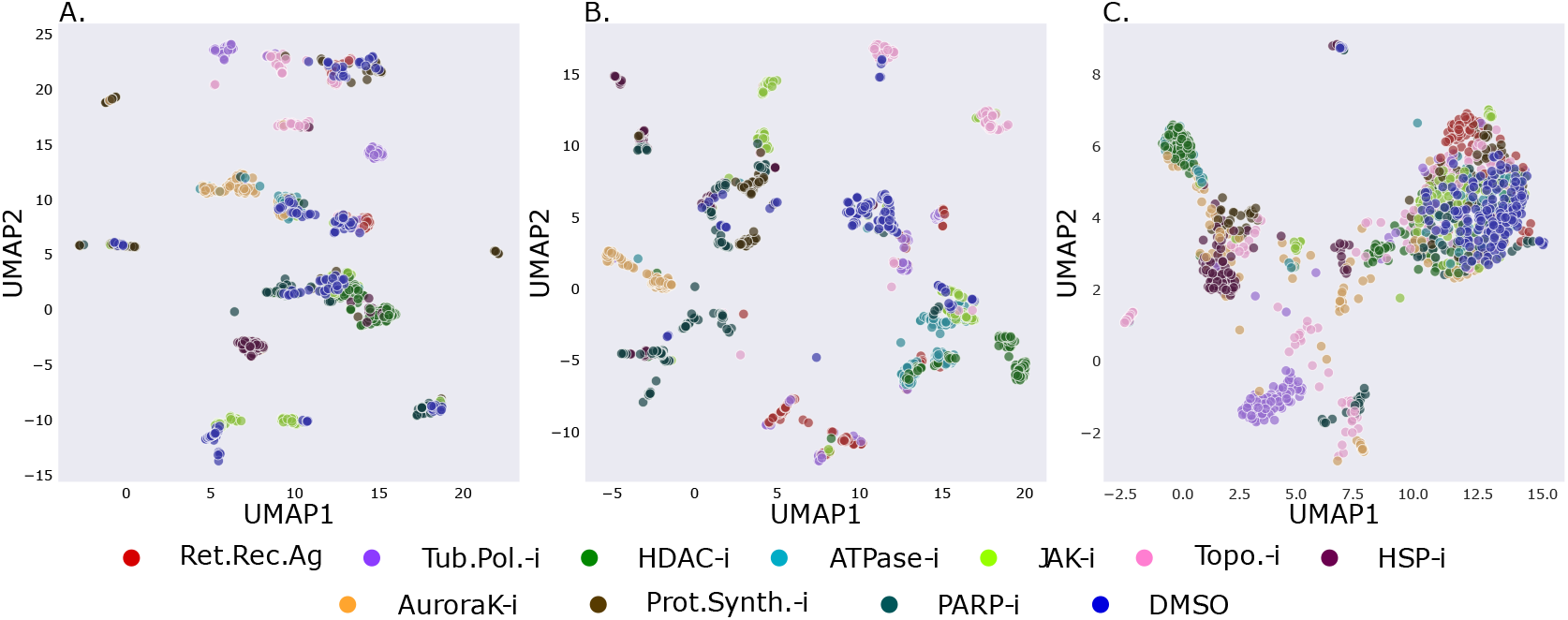
UMAP plots of features learned by the BF and FL models, and the raw CP features for the best performing split (split 2). **A**. BF features; **B**. FL features; **C**. CP cell-based features.

### Compound-level accuracy analysis

To further explore the performance differences between the models, we compared the compound-level accuracies, as shown in Figure 3. The Pearson correlation coefficient between BF and FL was 0.758, whereas between BF and CP it was 0.833 which indicates that prediction errors made by the BF models are more correlated with the CP benchmark than with the FL. The correlation between FL and CP models was the highest, at 0.875, as expected since they are both based on FL images. Bottom-right and top-left sections of the plots (shown by the dashed boxes in Fig 3) are “interesting” regions as they highlight the compounds where the BF performance was better, or worse than the counterparts, respectively. We identified seven compounds where the BF performance was consistently better than both FL and CP, which suggests that there may be cellular compartments or fine details picked up in the BF images, useful for MoA prediction, that are not detectable based on the Cell Painting protocol. In contrast, there was only one compound for which the BF performance was consistently worse than both the FL and CP models. The results indicate that there are multiple compounds in the dataset that exhibit morphological changes which can be picked up by the deep learning models applied to BF images.

**Figure 3:**
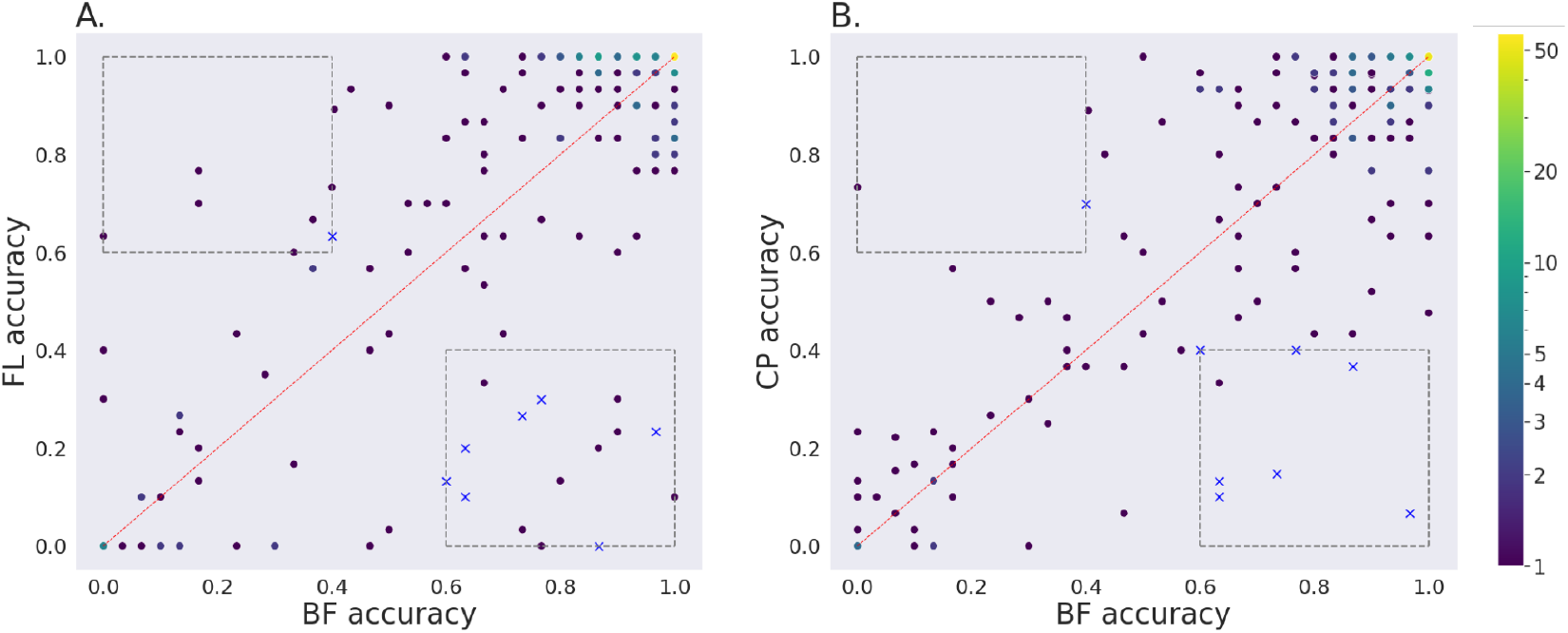
Comparison of the accuracy at the compound level, across all five test sets, for the BF models with respect to FL models, and CP feature-based models. Each dark dot represents a compound. Brighter dots represent multiple compounds with the same accuracy score. **A**. BF against FL; **B**. BF against CP. In the boxes at the bottom right and top left, thresholded at accuracy values of 0.6 and 0.4, the compounds shown with blue crosses were consistently better for BF than both FL and CP or consistently worse, respectively.

### Grit score analysis

For assessing the reproducibility of a compound treatment and its perturbation strength (morphological difference) relative to a control (in our case the DMSO) one can compute a grit score (https://github.com/cytomining/cytominer-eval, [14]). Based on CP features (extracted for the nuclei, cytoplasm and the entire cells in the FL images) we computed the grit scores for all the imaging sites used. A grit score of three for an imaging site means that on average the site is three standard deviations more similar to replicate sites for the same compound than it is to DMSO controls. The grit scores for some compounds (5 out of 231 compounds) were not calculated due to the images failing quality control, such as no cells present and out-of-focus images, and hence were not included in the grit-based analysis.

Figure 5 in the Supplementary section shows the counts of correct and incorrect classifications relative to the grit score for the three cases. For BF models, the error rate is higher for the lower grit scores and lower as the grit score becomes higher. The Area Under the ROC curve (AUROC) scores for logistic regression models (site-level predictions against the grit scores) were 0.662 for BF, 0.578 for FL, and 0.627 for CP. Hence, the grit score can predict the accuracy of the BF models better than the FL and CP-based models. This suggests that highly discriminative morphological features are present in the BF images of the compounds with higher grit scores, which can be leveraged to predict the MoAs with higher accuracy.

The grit scores for “interesting” cases in Fig 3 was also examined. The grit score for the seven compounds where BF models outperform both FL and CP models was in the range of 3.04-6.33 (4.61*±*1.16), and the grit score for the compound where BF accuracy was consistently worse was 0.550.

From the experiments in the previous sections the following observations can be made:

- BF models have difficulty in predicting DMSO (Table 1(b)) and DMSO features are not well separated from the other MoAs, as shown in Figure 2C.
- The FL models perform comparably to the CP models, as evident in Table 1(a).
- The BF models make more errors than the FL models for samples with lower grit scores, as shown in Figure 4 (dashed blue line).

**Figure 4:**
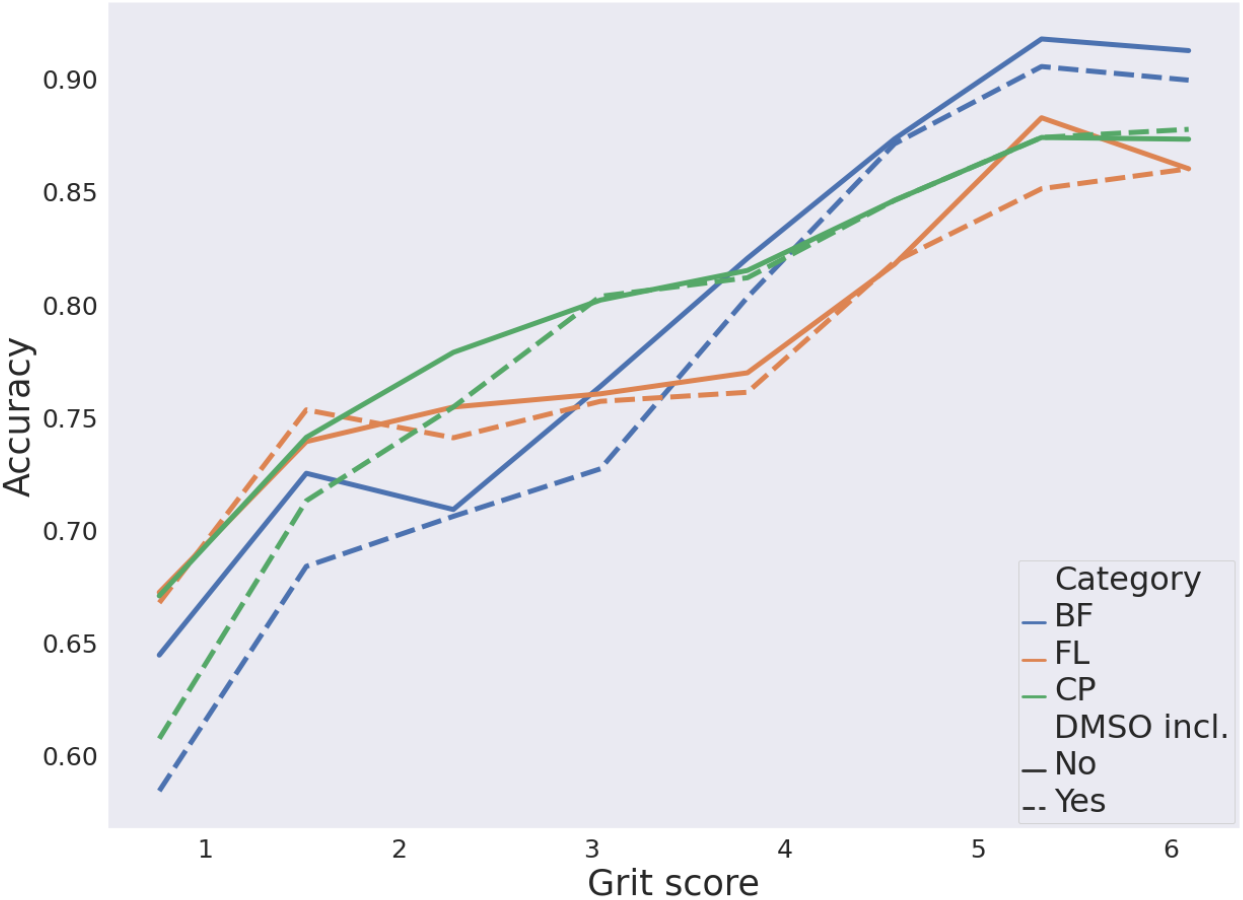
Comparison of accuracy at different grit scores for BF (blue), FL (orange), and CP feature-based (green) models across all five test sets. The dotted lines represent the score when DMSO was included in the experiments and solid lines represent the results when DMSO was excluded.

Based on these observations we hypothesized that in the absence of a clear separation between DMSO and some of the MoA classes, the DMSO samples assist the FL models in learning the subtle intensity difference between compounds with lower grit scores. However, in the absence of such intensity features in the BF data, the models face difficulties in separating the compounds from the DMSO and learning MoA-specific features. To test this hypothesis, we explored the effects of removing the DMSO class from the models.

### Models without DMSO as a predictive class

With the exclusion of the DMSO class, the BF models actually outperformed both the FL and CP models (see Table 2). The average performance of BF models increased by approx. 6%, outperforming the FL models by approx. 2% and performing equivalently to the CP feature-based models. This suggests that subtle morphological changes relevant for MoA prediction are present in the BF data, and by excluding DMSO, the BF models can learn these features and outperform the FL models.

**Table 2:** Comparison of results from the models trained on BF and FL images and CP features when DMSO was excluded as a predictive class.

(a) Macro-F1 scores on the test sets across the five data splits for BF, FL and CP models.
| Modality | BF | FL | CP |
| --- | --- | --- | --- |
| Split 1 | 0.765 | 0.753 | **0.781** |
| Split 2 | 0.821 | 0.809 | **0.829** |
| Split 3 | 0.767 | **0.782** | 0.775 |
| Split 4 | **0.746** | 0.714 | 0.729 |
| Split 5 | **0.795** | 0.718 | 0.763 |

(b) F1 scores per MoA, across all five test sets, for BF, FL, and CP models.
| MoA | BF | FL | CP |
| --- | --- | --- | --- |
| ATPase-i | **0.760** | 0.706 | 0.731 |
| AuroraK-i | **0.744** | 0.743 | 0.700 |
| HDAC-i | **0.800** | 0.785 | 0.772 |
| HSP-i | **0.744** | 0.708 | 0.716 |
| JAK-i | **0.756** | 0.671 | 0.705 |
| PARP-i | 0.890 | 0.878 | **0.960** |
| Prot.Synth.-i | **0.884** | 0.767 | 0.799 |
| Ret.Rec.Ag | 0.682 | 0.765 | **0.815** |
| Topo.-i | 0.733 | 0.715 | **0.771** |
| Tub.Pol.-i | **0.891** | 0.852 | 0.871 |
| Macro average | **0.788** | 0.759 | 0.784 |

This effect is also indicated in Figure 4 where the accuracy of the models is compared against the grit scores. We considered the grit score range of 0-6 as higher grit scores can result from cases where the dosage of the compound was too high and became cytotoxic to the cells. The BF accuracy increased by approximately 5% at the lower grit scores (0-2) when DMSO was excluded. An unexpected discovery can be seen in Figure 4 where the BF accuracy at higher grit scores (4-6) is higher than the FL and CP models, both with and without DMSO. This suggests that in FL images the morphological feature changes occurring at higher grit scores are not learned by FL models and they are also not represented by the extracted CP features. This could mean either that they are harder to learn or extract from FL images (e.g. signal saturation or channel bleed-through?), or that they are not present in the Cell Painting images. For comparative purposes, Figure 6 in the Supplementary section shows the counts of correct and incorrect classifications relative to the grit score when DMSO was excluded as a predictive class.

## 4 Conclusions and Future work

In our work, we found comparable predictive performance for models based on BF images to those based on both FL images and CP features derived from them. We found that the BF models made similar prediction errors to both the FL and CP models, although more so with the CP models. The BF models struggled to predict DMSO for cases where it was not well separated from some of the MoAs, however, their performance improved and they outperform the FL models when DMSO was not included as a predictive class. This shows that the features required to delineate the MoAs are present in the BF images and can be extracted by deep learning models.

Had the performance been significantly poorer for the BF models, than those based on FL, one avenue for exploration would have been to perform virtual staining [15, 16, 17] to generate virtually stained images from which to subsequently base the MoA prediction. However, going via this route would lose any morphological information that is only present in the BF images. We had several cases of compounds that were considerably better predicted by the BF model, relative to both the FL and CP models, suggesting that there are cellular compartments/organelles potentially picked up in the BF images that are not stained for in the Cell Painting protocol (such as membraneous organelles – lysosomes, endosomes and peroxisomes). We plan to investigate these cases further alongside those compounds for which the opposite was true (i.e. where the FL models performed well but the BF models did not). We hypothesize that this latter case may be a result of cellular compartments stained for in the Cell Painting protocol, which were useful for prediction, but which are barely visible or not accessible in the BF images (such as the f-actin cytoskeleton).

The fact that deep learning can be used on BF images for MoA prediction holds great promise for time-lapse studies, for which using FL data is problematic. With trained BF models we can track the cell populations over time to explore how the features evolve towards the MoAs after drug administration and how quick the process is. A wealth of interesting information will likely come from simply visualizing these temporal dynamics.

## Acknowledgments

The work was supported by the Swedish Foundation for Strategic Research (grant BD15-0008SB16-0046) and the European Research Council (grant ERC-2015-CoG 683810). The computations were enabled by the supercomputing resource Berzelius provided by National Supercomputer Centre at Linköping University and the Knut and Alice Wallenberg foundation. We also thank Maris Lapins for providing the Grit scores.

## 5 Supplementary

### 5.1 Data acquisition

#### Cell culture

The human osteosarcoma cell line U2OS (ATCC; HTB-96) was cultured in Dulbecco’s Minimum Essential Media (Gibco cat. no. 31885023) supplemented with 10% (v/v) fetal bovine serum (Gibco cat. no. 10500064), and 100 U/ml penicillin, and 100 μg/ml streptomycin (Gibco cat. no. 15140122). Cells were kept in a 37 °C humidified incubator with 5% CO2 atmosphere. We confirmed that the U2OS cell line was free from mycoplasma using the luminescence-based MycoAlert kit (Lonza cat. no. LT07-218).

#### Compounds

Compound handling was performed by the SciLifeLab Compound Center (CBCS, Solna, Stockholm). In brief, chemicals were solubilized in DMSO at a concentration of 10mM, then 40nl of each compound was dispersed using the Echo liquid handler into Falcon optilux microplates (Falcon, cat. no. BD353962) and stored at -20 °C prior to experimentation. Compounds were distributed over the plates with three technical replicates and two biological replicates. To reduce bias by positional effects in the microwell plates, the conditions were distributed over the plates using PLAID (Plate Layouts using Artificial Intelligence Design, https://github.com/pharmbio/plaid).

#### Cell Painting

The Cell Painting protocol [5] was followed with a few adjustments. A Biotek MultiFlo FX was used for dispensing cells and solutions and a Biotek 405 LS microplate washer was used for washing steps. (A robotic arm (UR3) moves plates between the incubator, plate hotel, and washer and dispenser, using tailored software for scheduling (more info: https://github.com/pharmbio/aros). In short, 40 *μ*l of cells were dispensed on top of DMSO solubilized compounds at a density of 1100 cells/well. The plates were incubated for 48 hr at 37 °C at 5% CO2 atmosphere. Then, assay plates were washed with 80 *μ*l 1× PBS (Thermo Fisher, cat.no 11510546), followed by addition of 30 *μ*l MitoTracker (Invitrogen; M22426) in prewarmed Live Cell Imaging solution. After 20 minutes of incubation, the cells were washed with 80 *μ*l 1× PBS and fixed in 80 *μ*l of 4% PFA (Histolab; 02176) for 20 min. The plates were washed three times, followed by permeabilization with 80 *μ*l of 0.1% Triton X-100 for 20 min at room temperature and washed three times with PBS. Then, 20 *μ*l staining mixture was added to each well reaching a final well-concentration of 1 *μ*g/ml Hoechst, 15 *μ*g/ml Wheat germ agglutinin, 10 *μ*l/ml Phalloidin, 4 *μ*M SYTO 14 and 80 *μ*g/ml Concanavalin A, and was incubated for 20 min. The targets for each of the stains were: DNA (Hoechst); mitochondria (MitoTracker); Golgi apparatus and plasma membrane (Wheat Germ Agglutinin); F-actin (Phalloidin); nucleoli and cytoplasmic RNA (SYTO 14); and the endoplasmic reticulum (Concanavalin A/Alexa Fluor 488). Stains were removed and plates were washed three times with 1X PBS prior to imaging.

#### Image acquisition

Microplates were imaged using a widefield high-throughput ImageXpress Micro XLS (Molecular Devices) microscope with a 20X objective with laser-based autofocus. Fluorescent images were captured using five fluorescent channels. Excitation spectra were set to 377/50 nm (Hoechst), 628/40 nm (Mitotracker), 562/40 nm (Phalloidin and Wheat germ agglutinin), 531/40 nm (SYTO 14) and 482/35 nm (Concanavalin A). Emission filters were set to detect signals between 447/60 nm (Hoechst), 692/40 nm (MitoTracker), 624/40 nm (Wheat Germ Agglutinin and Phalloidin), 593/40 nm (SYTO 14) and 536/35 (Concanavalin A). For each well, a total of nine sites of view were captured using a single z-plane targeting the cell compartment of interest. Brightfield images were captured under transmitted light, for five fields of views and 6 focal planes, 2*μ*m apart from each other. The Hoechst staining was used for autofocus. Images were saved as 16-bit grayscale TIFF files without binning (1996×1996 pixels).

### 5.2 Model training

During training, the models were optimized with stochastic gradient descent (SGD), with a momentum of 0.9 and weight decay of 1e-4. The models were trained for 750 epochs with a batch size of 64. A linear warmup cosine annealing learning rate scheduler was applied, whereby the learning rate was increased linearly from 0.005 to 0.05 for the first 10 epochs and then reduced with cosine annealing to 5e-5 at the end. Random cropping (1024×1024 pixels) followed by resizing (to 512×512 pixels), grid shuffle, flipping, and 90-degree rotation augmentations were also used. All the experiments were run with a fixed random seed and all the hyperparameters were kept the same for both the BF and FL models.

### 5.3 Normalization

For the BF data, we explored performing normalization at the image and well level, thus removing the relative intensity differences at the image and well-level, respectively. Additionally, we explored standardizing at the plate level, using the data from all the treatments and DMSO wells on each plate, and finally applying the same method as used for the FL data, i.e. DMSO plate-level normalization.

**Table 3:** Macro-F1 scores on the test sets across the five data splits with different normalization techniques applied to the brightfield (BF) data and DMSO normalization applied to the fluorescence (FL) data.

| Modality | BF |  |  |  | FL |
| --- | --- | --- | --- | --- | --- |
|  | Well Norm | Image Norm | Plate Norm | DMSO Norm | DMSO Norm |
| Split 1 | 0.560 | 0.598 | 0.651 | <u>0.688</u> | **0.777** |
| Split 2 | 0.639 | 0.718 | 0.733 | <u>0.773</u> | **0.799** |
| Split 3 | 0.562 | 0.607 | 0.663 | <u>0.700</u> | **0.793** |
| Split 4 | 0.549 | 0.633 | 0.682 | <u>0.731</u> | **0.738** |
| Split 5 | 0.577 | 0.643 | 0.685 | <u>0.701</u> | **0.716** |

### 5.4 Grit score comparison

**Figure 5:**
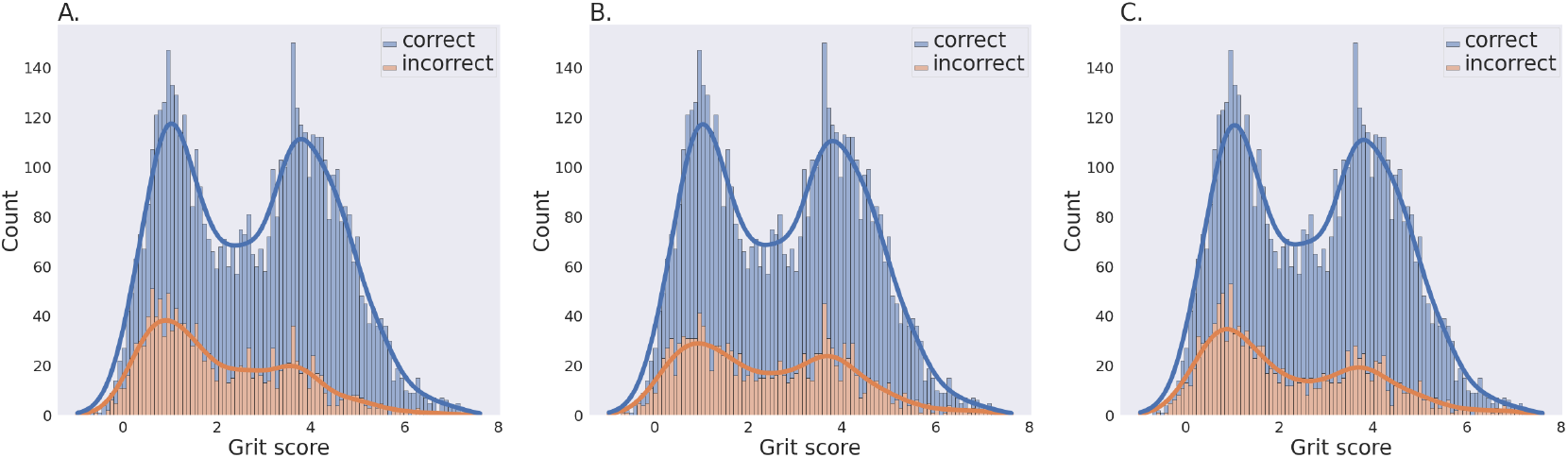
Counts of correct and incorrect classifications relative to the grit score, across all five test sets, at the imaging site level for: **A**. BF models; **B**. FL models; and **C**. CP feature-based models.

**Figure 6:**
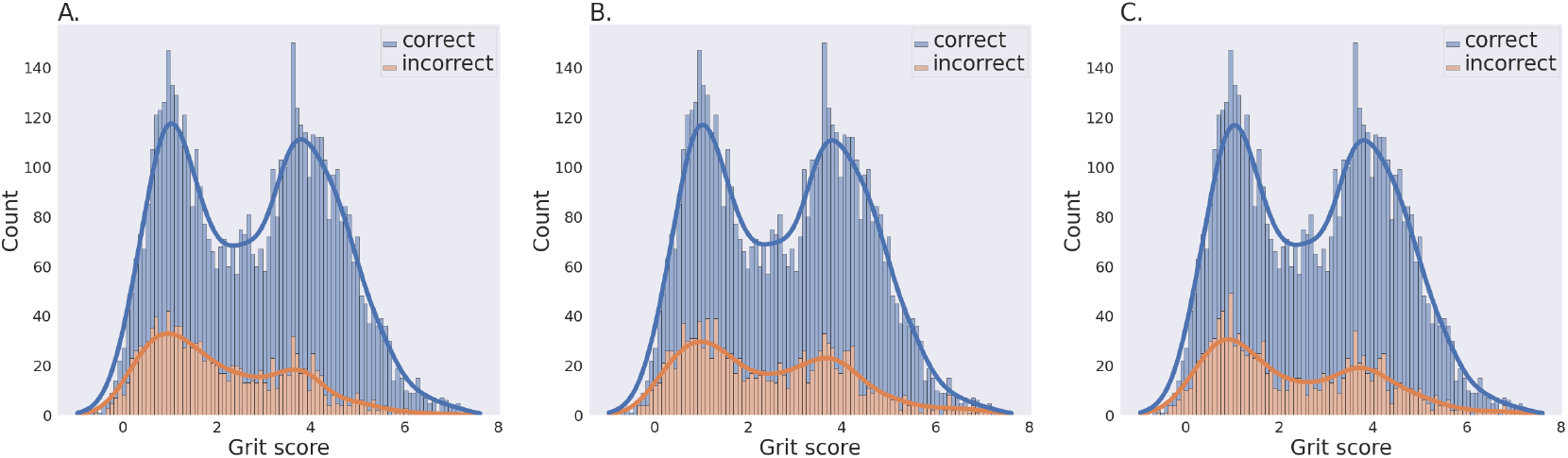
Counts of correct and incorrect classifications relative to the grit score (when DMSO is excluded as a predictive class), across all five test sets, at the imaging site level for: **A**. BF models; **B**. FL models; and **C**. CP feature-based models.

### 5.5 Compound-level accuracies with the DMSO class excluded

**Figure 7:**
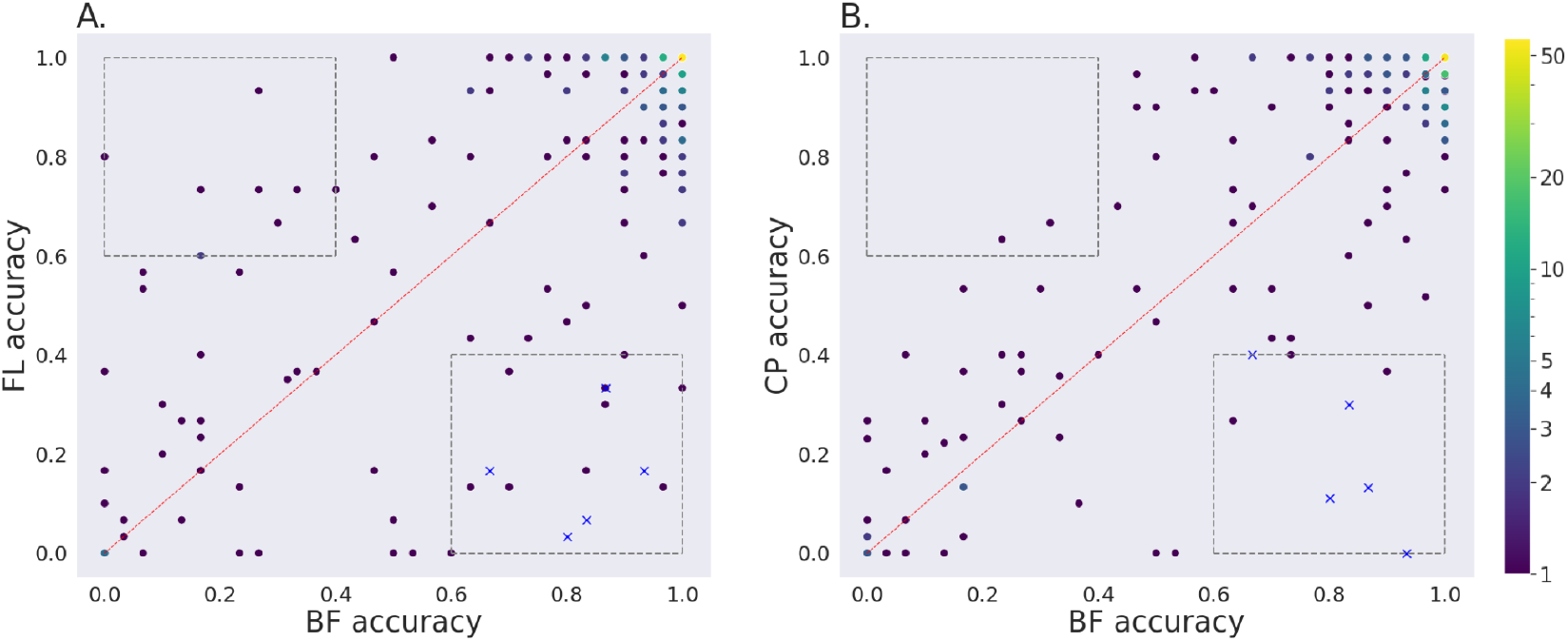
Comparison of the accuracy at the compound level with the DMSO class excluded, across all five test sets, for the BF models with respect to FL models, and CP feature-based models. Each dark dot represents a compound. Brighter dots represent multiple compounds with the same accuracy score. **A**. BF against FL; **B**. BF against CP. In the boxes at the bottom right and top left, thresholded at accuracy values of 0.6 and 0.4, the compounds shown with blue crosses were consistently better for BF than both FL and CP or consistently worse, respectively.

